# Controlling pallidal oscillations in real-time in Parkinson’s disease using evoked interference deep brain stimulation (eiDBS): proof of concept in the human

**DOI:** 10.1101/2021.05.22.445251

**Authors:** David Escobar Sanabria, Joshua E. Aman, Valentina Zapata Amaya, Luke A. Johnson, Hafsa Farooqi, Jing Wang, Meghan Hill, Remi Patriat, Kelly Sovell-Brown, Gregory F. Molnar, David Darrow, Robert McGovern, Scott E. Cooper, Noam Harel, Colum D. MacKinnon, Michael C. Park, Jerrold L. Vitek

## Abstract

Approaches to control basal ganglia neural activity in real-time are needed to clarify the causal role of 8-35 Hz (“beta band”) oscillatory dynamics in the manifestation of Parkinson’s disease (PD) motor signs. Here, we show that resonant beta oscillations evoked by electrical pulse with precise amplitude and timing can be used to predictably suppress or amplify spontaneous beta band activity in the internal segment of the globus pallidus (GPi) in the human. Using this approach, referred to as closed-loop evoked interference deep brain stimulation (eiDBS), we could suppress or amplify frequency-specific (16-22 Hz) neural activity in a PD patient. Our results highlight the utility of eiDBS to characterize the role of oscillatory dynamics in PD and other brain conditions, and to develop personalized neuromodulation systems.

## Introduction

While much research has been dedicated to understanding the pathophysiology of Parkinson’s disease (PD), the neural circuit dynamics underlying the manifestation of specific motor signs remain to be demonstrated. Current theories propose that the amplitude and incidence of 8-35 Hz “beta” band oscillations, synchronized throughout the basal ganglia thalamocortical (BGTC) circuit, are associated with the severity of motor signs[1–6]. Although changes in bradykinesia related to levodopa and deep brain stimulation (DBS) treatments have been shown to correlate with the power of local field potential (LFP) activity in the subthalamic nucleus (STN) and internal segment of the globus pallidus (GPi) [1–10], no study has deductively or conclusively demonstrated their causal relationship. DBS yields therapeutic benefit via continuous delivery of high-frequency (~130 Hz) electrical pulses in the STN or GPi. DBS can also suppress beta band oscillations in the target while improving motor function. Yet, the mechanisms by which high-frequency stimulation produces these therapeutic and physiological effects are not clear. This knowledge gap limits our ability to assess the causal relationship between suppression of oscillations attained with DBS and the improvement of motor signs. Does high-frequency stimulation directly reduce beta band oscillations and thereby produce a therapeutic effect? Or is the reduction in beta band oscillations during DBS secondary (or unrelated) to the therapeutic effects of DBS? Previous studies have attempted to answer these questions using 20 Hz electrical stimulation of the STN, with the idea that stimulation at this frequency may promote the generation of STN rhythms in the beta band. While studies have reported that 20 Hz STN stimulation can worsen bradykinesia in some PD patients[11–13], others have challenged this idea and have shown no effect[14,15]. The aforementioned studies highlight the need for approaches that can control beta band oscillations in real-time, without employing high-frequency stimulation, to characterize the role of these oscillations in the manifestation of PD.

In the current study, we demonstrate that resonant oscillations in the human GPi evoked by stimulation pulses in the GPi can be employed to suppress or amplify frequency-specific spontaneous GPi oscillations in real-time without utilizing high-frequency stimulation. We used a feedback (closed-loop) control strategy in which stimulation pulses were delivered in the GPi with precise amplitude and timing relative to the targeted GPi oscillations to evoke neural responses that suppress or amplify these oscillations. The rationale behind this approach, referred to as closed-loop evoked interference DBS (eiDBS), is that synaptic-related neural responses evoked by electrical pulses can “override” spontaneous, synaptic-related oscillations via synaptic integration when the pulses are delivered with precise amplitude and timing relative to the phase of spontaneous oscillatory activity [16]. See schematic of eiDBS in Fig. 1A.

**Figure 1.**
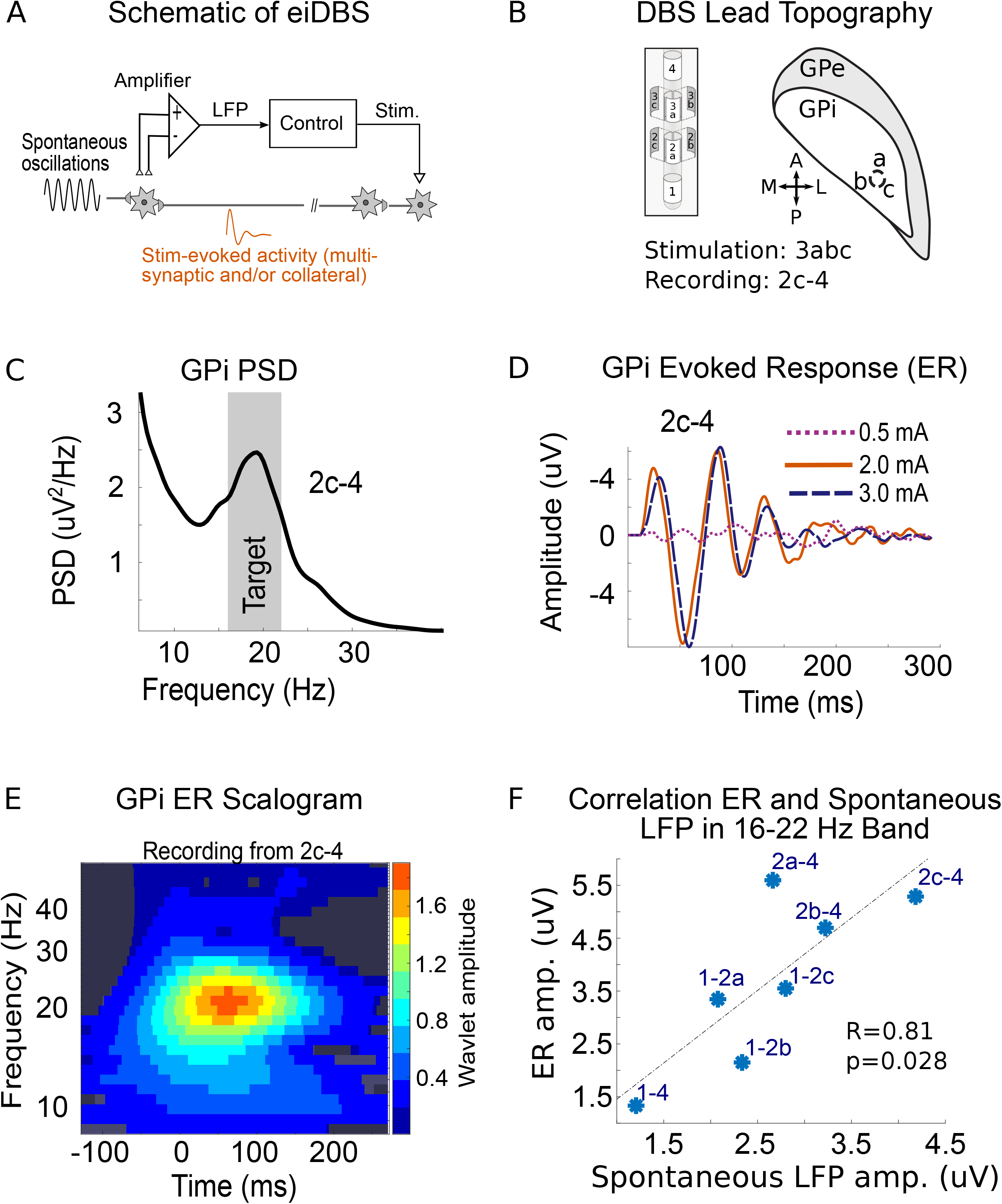
(A) Schematic of closed-loop evoked interference DBS. eiDBS delivers stimulation pulses with precise amplitude and timing to evoke resonant neural responses that overrides spontaneous oscillations via constructive or destructive interference. (B) Diagram of directional DBS lead and localization of directional contacts in the GPi adapted from [22]. The 2D slice in the axial plane is at the depth of ring 3 in the DBS lead. The orthogonal coordinate frame depicted on the axial plane consists of the anterior (A)-posterior (P) and medial (M)-lateral (L) axes. (C) Power spectral density (PSD) of LFPs recorded from contacts 2c-4. (D) Neural responses in the GPi evoked by stimulation in the GPi with currents equal to 0.5, 2.0, and 3.0 mA. (E) Wavelet transform scalogram (time-frequency map) of GPi response evoked by 3 mA stimulation pulses. Regions where the ER is not significantly greater in amplitude than surrogate data are depicted in gray, predominately in the upper right and left hand corners of the scalorgram. Colored regions (not gray) correspond to regions where the ER is significantly greater than surrogate data (p<0.01). (F) Scatter plot of ER amplitude vs. spontaneous activity amplitude in the 16-22 Hz band as observed across differential recordings from the DBS directional lead. Scalar measures of ER amplitude for each montage are equal to the sum of scalogram values over frequencies in the targeted band (16-22 Hz) at the time where the maximum ER amplitude is observed. Scalar measures of the spontaneous LFP amplitude are equal to the sum of amplitude spectral density (square root of PSD) values over frequencies in the targeted band. R=0.8 is the correlation coefficient associated with the ER and spontaneous activity data points.

We tested eiDBS in a PD patient implanted with a directional DBS lead in the GPi and evaluated the suppression and amplification capabilities of this neuromodulation approach. eiDBS was capable of suppressing or amplifying GPi oscillations in the targeted frequency band (16-22 Hz) in real-time. Stimulation-evoked responses (ERs) that mediated this modulation resonated in the beta band, within the same frequency range where the peak power of the spontaneous LFPs was located. Because the ERs resided in the beta band, eiDBS required less stimulation amplitude to modulate beta oscillations than the stimulation needed to modulate neural activity in other frequency bands. This study provides the rationale for future studies to assess the causal role (direct or indirect) of oscillatory dynamics in PD using eiDBS. It also highlights the prospect of developing personalized neuromodulation systems based on interference between stimulation-evoked and spontaneous neural activity.

## Material and Methods

### Patient and surgical procedure

All patient procedures were approved by the University of Minnesota Institutional Review Board (IRB protocol #1701M04144) with consent obtained according to the Declaration of Helsinki. This study was conducted with a male patient (55 years old) diagnosed with idiopathic PD ~6 years before unilateral (right side) GPi DBS surgery. Intraoperative microelectrode mapping was used to identify the sensorimotor region of GPi for DBS lead placement[17–20]. Following intraoperative microelectrode mapping, a directional DBS lead (Boston Scientific Vercise Cartesia model DB-2202-45; 1.5 mm contact height with 0.5 mm vertical spacing) was implanted. See diagram of the DBS lead in Fig. 1B. After DBS implantation, a lead extension was tunneled to a subcutaneous pocket in the chest and connected to another extension that was externalized through an abdominal incision[21]. Five days later, the patient was admitted at the University of Minnesota Clinical Research Unit, where experiments took place over the course of two days.

### Electrode localization

The location of the DBS lead contacts in the GPi was confirmed based on information obtained during intraoperative electrophysiological mapping as well as co-registered preoperative 3T MRI and postoperative CT scans[21]. Electrode localization and orientation for this patient was previously described in [22]. Briefly, the lead orientation relative to the brain was derived using a modified version of the DiODe algorithm[23] and based on unique artifacts of the lead contacts and fiducial marker superior to the most distal contact from the lead tip. The orientation of the lead was confirmed with information extracted from fluoroscopy and X-ray images acquired intraoperatively.

### Postoperative externalized recordings

The externalized lead extension was connected to an ATLAS neurophysiological recording system (Neuralynx, Bozeman, MT, USA) via customized connectors. Reference and ground EEG electrodes were placed on the scalp along the midline. An FDA-approved neurostimulator (g.Estim, g.tec, Schiedlberg, Austria) was employed to deliver current-controlled stimulation. A skin-surface patch electrode was placed on the chest contralateral to the DBS lead to serve as a current return for monopolar electrical stimulation. On day 1 of the externalization recordings, we collected resting LFP data in the off-stimulation condition and during low-frequency (3 Hz), monopolar stimulation (0.5, 2, and 3mA with 60 us pulse width). These data were used to characterize evoked responses and calculate eiDBS parameters that maximize suppression or amplification of beta band oscillations. See the **Optimization of eiDBS** Section. In the morning of day 2, we tested eiDBS with the parameters found on day 1 in the off-medication state, 16 hours after the last dose of carbidopa/levodopa (1 pill, 25/100 mg).

### Characterization of stimulation-evoked responses

All data analyses were performed using custom software developed in MATLAB (The MathWorks, Natick, MA). Evoked responses in the GPi were computed by averaging LFP segments aligned with stimulation pulses. Stimulation artifacts were removed as described below in the **eiDBS implementation** Section. The amplitude of evoked responses in the frequency and time domain were characterized using spectrograms (time-frequency maps) calculated with the wavelet transform. We assessed whether observed evoked responses in the spectrograms were the result of chance using a permutation test without replacement, performed by randomly sampling 10,000 resting state LFP segments and computing scalograms for each permutation. By using the permutation distribution of surrogate scalograms, we computed the p-value of the original wavelet value at each frequency and time. A false discovery rate (FDR) correction for multiple tests in the time and frequency domains was applied to the p-values. Values of wavelet amplitudes with corrected p<0.01 were considered significant.

The effect of stimulation pulses on the ER temporal dynamics was characterized using a saturation nonlinearity (static) connected to a system of linear differential equations (dynamics) as depicted in Fig. 2A. This model has been shown to estimate the response of ER to stimulation pulses accurately[16]. The parameters of the differential equations were obtained using the instrumental variable system identification approach[24]. A transfer function with four poles and one zero was selected using Akaike’s Information Criterion (AIC) to minimize both the model prediction error and the number of estimated parameters[24]. The differential equations describing the ER dynamics are presented in Appendix A. The largest gain of the ER input-output transfer function is at 19.9 Hz (largest gain in the Bode magnitude diagram). Therefore, periodic stimuli at this frequency leads to the highest amplitude evoked responses, implying that a minimum stimulation amplitude is needed to suppress or amplify oscillations at 19.9 Hz via eiDBS as compared to oscillations at other frequencies.

**Figure 2.**
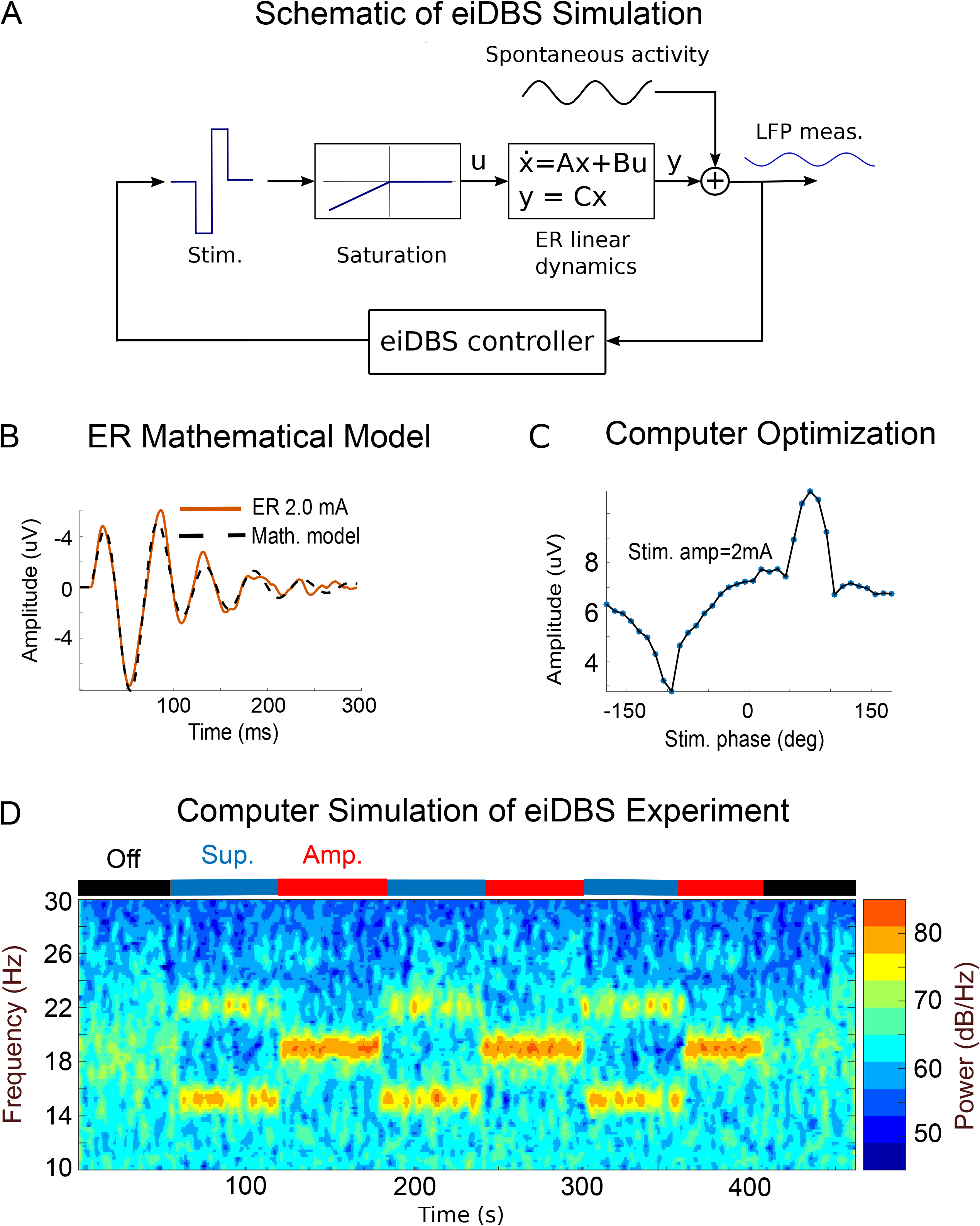
(A) Schematic of eiDBS computer simulation in which stimulation pulses are triggered by a closed-loop controller based on real-time LFP measurements. The model consists of a static saturation nonlinearity that allows us to capture the symmetric response of the GPi to cathodal and anodal stimulation pulses. The negative phase of the stimulus is the input of a linear time-invariant system of differential equations that reproduce the evoked response temporal dynamics. The LFP measurement is modeled as the linear superposition of the ER and spontaneous oscillations. (B) Measurement and mathematical model of GPi ER for a stimulation amplitude of 2mA. (C) Effect of eiDBS (2 mA) delivered at different phase angles on the mean amplitude of targeted neural activity in the 16-22 Hz band (computed over 60 s). The curve indicates that the optimal phase angles for suppressing and amplifying the targeted neural activity are −85 and 95 deg, respectively. (D) Spectrogram (LFP power in the time-frequency domain) of computer simulation in which eiDBS was delivered with parameters found to maximize the suppression and amplification of 16-22 Hz oscillations.

### Real-time neural control infrastructure

An industrial computer dedicated to modulating brain activity in real-time with a closed-loop delay of less than 1ms was used to implement eiDBS (Mobile Target Machine, SpeedGoat, Bern Switzerland). The real-time computer was connected to the ATLAS recording system via a fiber optic link (Gigabit Ethernet) to read LFP data at 24 KSamples/s and to a g.Estim stimulator via a digital interface to control individual stimulation pulses.

### Implementation of eiDBS

The algorithm used to implement eiDBS has been previously described in preclinical studies with nonhuman primates[16]. Briefly, this algorithm consists of 1) acquiring LFP data from the ATLAS system at 24 Kilosamples/s, 2) suppressing stimulation artifacts using a dynamic template model of the artifact and blanking residual artifacts (2.6 ms blanking duration), 3) filtering differential LFPs in the targeted frequency band using a 2^nd^ order Butterworth filter and down-sampling LFPs at 3 KHz, 4) computing the instantaneous phase and amplitude of the filtered LFP using a Hilbert transformer filter, and 5) triggering single stimulation pulses at specific phases of the neural oscillations. Stimulation was not delivered if the amplitude envelope of these oscillations was less than a prescribed threshold. This threshold was equal to the 20th percentile of the oscillations’ amplitude envelope, calculated in the off-stimulation condition. Because we sample the data and remove the artifact at 24 Kilosamples/second and have amplifiers with a large input range (+/-132 mV), we can consistently and accurately capture the shape of the artifacts and robustly remove the artifacts in real-time. Ringing effects associated with filtering are not observed in our setup because we remove the artifacts before the LFPs are low-pass filtered and down-sampled.

For sensing, we selected contacts 2c and 4, whose differential LFP had the largest signal-to-noise ratio in the frequency band targeted for modulation (16-22 Hz). This frequency band was centered on the peak of the LFP power spectral density. The sensing montage was selected among the following combinations: 1-3a, 1-3b, 1-3c, 2a-4, 2b-4, and 2c-4. We assessed these electrode combinations to enable monopolar stimulation to be delivered through the ring located between sensing electrodes and thereby minimize stimulation artifacts via differentiation of the sensing electrodes. Stimulation was delivered using monopolar stimulation through ring 3abc (segments 3a, 3b, and 3c tied together). Monopolar stimulation was utilized to minimize the size of electrical artifacts via differentiation of potentials measured in electrodes equidistant to the monopole. One should not that artifact asymmetries are always observed in the recordings, even when computing differential potentials between electrodes with the same surface area and the same distance to the monopolar current source. Therefore, the artifact removal routine described above is necessary, in addition to differentiation, to effectively suppress artifacts.

### Optimization of eiDBS

A computer optimization (search) for eiDBS was performed based on the ER mathematical models described above to determine the stimulation parameters that maximize suppression of neural oscillations in the targeted band[16]. We created patient-specific computer simulations using these evoked response models and simulated the closed-loop algorithms with a range of stimulation phase angles for a given stimulation amplitude. We characterized stimulation evoked responses and created artifact models for currents equal to 0.5, 2.0, and 3.0 mA only. Among these three current levels, 2.0 mA was selected because it offered a modulation effect equivalent to that attained with 3 mA (see the size of evoked responses in Fig. 1D) but with less stimulation energy. A search across phase angles (−180 to 175 deg with a 5-deg. resolution) for a stimulation current of 2 mA was performed to calculate the phase that minimized the amplitude of neural activity in the targeted frequency band. The same stimulation amplitude determined for eiDBS-suppression was also used for eiDBS-amplification. The stimulation phase for eiDBS-amplification was determined using the search described above with the given stimulation amplitude.

### Assessment of oscillatory activity

Artifacts were removed from the LFP raw data as described above in the **Implementation of eiDBS** Section, and then the LFPs were down-sampled at 3KHz for processing. The power of the LFP recordings in the time and frequency domain was characterized using spectrograms and the Welch method. We measured the average power in a specific condition (e.g., eiDBS suppression) using PSD curves computed with the Welch method. We assessed whether the amplitude of neural oscillations in the targeted frequency band changed when eiDBS was delivered by using scalar measurements of the oscillations’ amplitude envelope. These scalar measurements were computed by filtering the artifact-suppressed data in the targeted band, calculating the magnitude of the analytic signal via the Hilbert transform, and averaging the amplitude envelope over nonoverlapping windows of three-second duration and separated by one second. This separation time is greater than the maximum time between effectively independent data, calculated across conditions (off stimulation, amplification, suppression) by using the autocorrelation function[16,25,26]. Pairwise differences between scalar measurements of the oscillations’ amplitude in two different conditions were assessed via the Wilcoxon rank-sum test. The p-values resulting from this test were corrected for the two comparisons via the Bonferroni method. We assumed that the difference between measurements in the two conditions was significant when p < 0.05. We evaluated effect sizes using the Cohen signed (non-parametric) test (‘U3’)[27].

## Results

### LFP and evoked response oscillations matched frequency and location

We characterized spontaneous and stimulation-evoked neural activity and evaluated the feasibility of delivering eiDBS in a PD patient implanted with a directional DBS lead in the sensorimotor region of the GPi (Fig. 1B). Resting state, off-stimulation LFPs recorded from the patient’s GPi exhibited elevated beta band oscillations with a peak frequency of 19 Hz in the power spectral density (PSD). See Fig. 1C. We selected the frequency band between 16 and 22 Hz for modulation via eiDBS. Open-loop, low-frequency electrical stimulation pulses (2.93 Hz, 60 us pulse width) evoked neural responses with highest power at 20.8 Hz, near the peak frequency of the spontaneous LFP PSD (i.e., 19 Hz). See Figs. 1C,E. The amplitude of the GPi ERs was insignificant for stimulation currents equal to 0.5 mA but clear and significant for currents equal to 2 and 3 mA. This nonlinear response to stimulation is a characteristic of neural stimulation-evoked responses[16].

For eiDBS to have a true modulatory effect on spontaneous neural activity, the ERs measured with the DBS lead need to be generated by the same neuronal population that give rise to the LFP spontaneous oscillations. We used the spatial distribution of electric potentials across differential recordings from the directional DBS lead to evaluate whether the ER and LFP spontaneous oscillations were generated by the same neural source. Across all available differential potentials from the directional DBS lead, the amplitude of the ERs was highly correlated with the amplitude of the spontaneous beta oscillations (Correlation coefficient R=0.8, slope =1.37, p=0.028). See Fig. 1F. This high correlation is a necessary condition for the neural sources (monopolar or dipolar) generating the ER and spontaneous oscillations to be in the same location according to the Poisson equation of electrostatics[28,29]. The solution to this equation implies that the spatial distribution of electric potentials generated by two neural sources should be the same if these neural sources are at the same location.

### eiDBS was capable of suppressing or amplifying frequency-specific GPi activity

We constructed mathematical models of the ERs based upon the patient’s data using system identification techniques (Fig. 2A,B) as described in more detail in the Methods Section and preclinical work published previously[16]. These ER models are described by linear differential equations and a saturation element that transforms the biphasic stimulation pulses to monophasic pulses responsible for the neural response. Using the patient-specific ER mathematical models and recorded LFP data, we constructed a computer simulation (Fig. 2A,B) to characterize the neural modulation attained with eiDBS and search for the stimulation parameters (amplitude and phase angle) that maximized suppression and amplification of neural activity in the targeted band (16-22 Hz). See optimization curve in Fig. 2C and computer simulation with optimized parameters in Fig. 2D. The parameter search was performed with stimulation amplitudes for which ER and stimulation-artifact models were available from the recorded data (0.5, 2.0, and 3.0 mA). The computer simulation (Fig. 2D) indicated that when neural activity was suppressed in the targeted frequency band, there was amplification of activity in adjacent frequencies (~15 Hz and ~23 Hz).

We tested eiDBS in-vivo with the study participant using the stimulation parameters found to be optimal (stimulation at 2 mA and phase angles equal to −85 and 95 deg for suppression and amplification). eiDBS was capable of suppressing or amplifying pallidal activity in the targeted frequency interval (16-22 Hz). See Figs. 3A-D. eiDBS (suppression or amplification) was well tolerated by the patient. During the suppression stage of the eiDBS experiment, the median LFP amplitude in the targeted band decreased from 4.59 to 2.74 uV (p= 6.12e-08 with rank-sum test, Cohen’s U3 effect size=1). During the amplification stage, the median amplitude of oscillations in the targeted band increased from 4.59 to 7.25 uV (p=4.04e-8 with rank-sum test, Cohen’s U3 effect size=1). Suppression of neural activity in the targeted band resulted in an increase in the median amplitude of the LFP in the 12-16 Hz band (3.63 uV in the off stimulation vs. 5.01 uV in the suppression condition, p=4.5e-7 with rank-sum test, Cohen’s U3 effect size=1). While an increase in power at ~23 Hz is observed in the suppression condition (spectrogram of Fig. 3A), our statistical analysis indicated that no significant changes occurred in the 22-26 Hz band.

**Figure 3.**
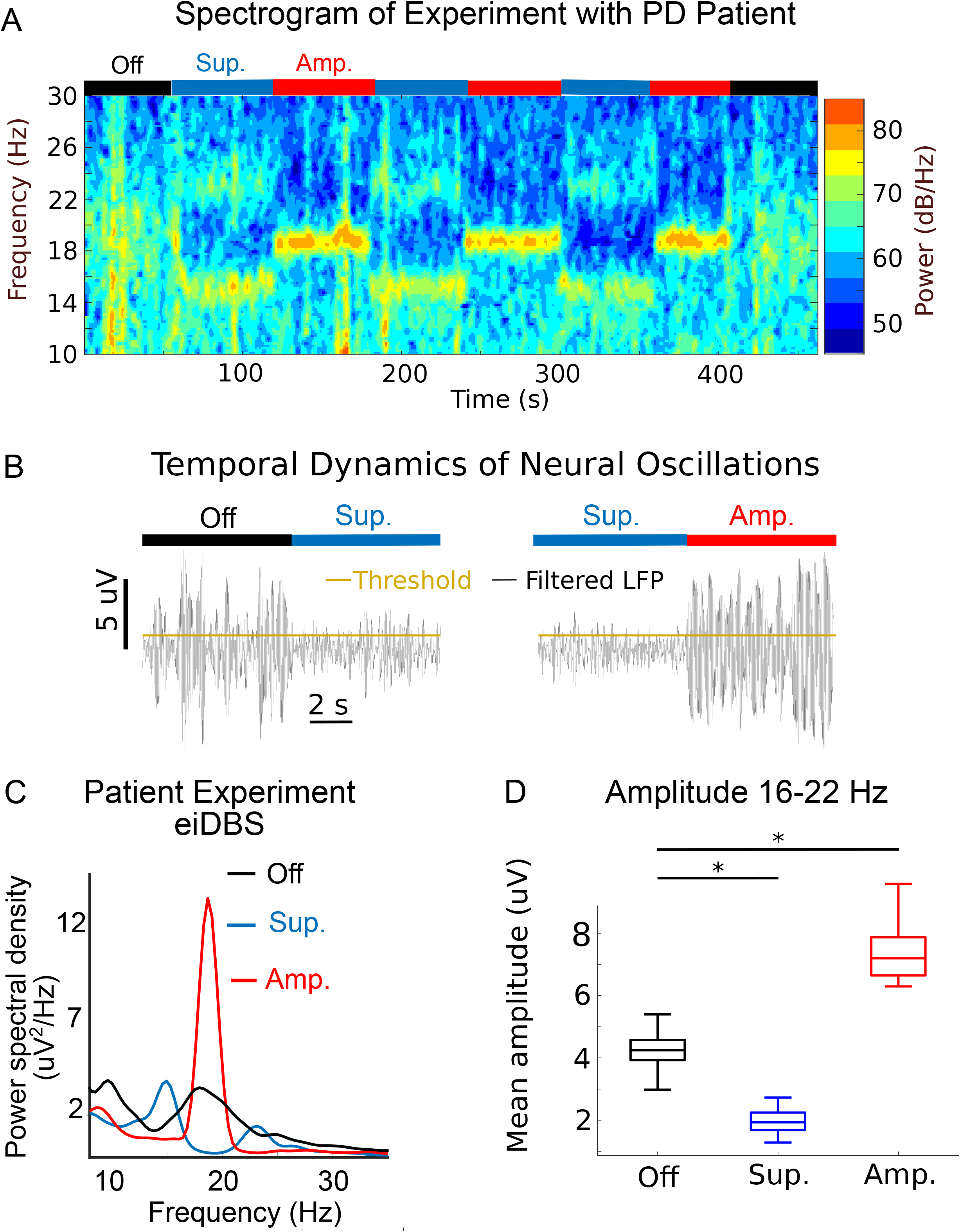
(A) Spectrogram (LFP power in the time-frequency domain) of LFP activity during periods in which eiDBS was delivered to suppress and amplify targeted GPi oscillations in a PD patient implanted with a directional DBS lead. (B) Temporal dynamics of modulated oscillations (filtered in the 17-21 Hz range for clear visualization of modulatory effects on ~19 Hz oscillations) in the off-stimulation to suppression, and suppression to amplification transitions. (C) Mean power spectral density (PSD) of LFP activity in the off-stimulation, suppression, and amplification conditions illustrates how eiDBS modulated spontaneous pallidal oscillations across frequencies. (D) Boxplot with independent measurements (mean over 3 s) of the oscillations’ amplitude envelope in the targeted band (16-22 Hz) in the off-stimulation, eiDBS-suppression, and eiDBS-amplification conditions. The box edges represent the interquartile range, and the horizontal line within each box represents the median. Most extreme data points not considered outliers are represented by the whiskers. The amplitude of oscillations significantly decreased from the off-stimulation to the suppression condition (p= 6.12e-08, Cohen’s U3 effect size=1) and increased from the off-stimulation to the amplification condition (p=4.04e-8, Cohen’s U3 effect size=1). P-values were corrected for the two comparisons made using the Bonferroni method. The symbol ⍰ indicates that the difference between conditions was statistically significant with the p-values listed above. The number of independent observations (mean amplitude over 3 s) used in this analysis was n=16 in the off-stimulation, n=37 in the suppression, and n=29 in the amplification condition.

### Stimulation-evoked responses in the GPi mediated the modulation achieved via eiDBS

The computer simulations constructed with the ER mathematical models and patient-specific LFP activity (Fig. 2D) predicted the modulatory effects of eiDBS in the suppression and amplification conditions during the in-vivo experiments with the subject. The ability of the computer simulation to predict the modulatory effect of eiDBS indicates that the ERs mediated the suppression or amplification of spontaneous oscillations in the patient, given that the ER mathematical model drove the computer simulations. Moreover, the ER transfer function (input-output dynamic map) indicates that the highest gain of this transfer function is at 19.9 Hz. See Methods Section. Therefore, periodic stimulation at 19.9 Hz yields evoked responses larger than stimulation at any other frequency. When these evoked responses have the same amplitude and are out of phase with spontaneous oscillations at the same frequency (i.e., eiDBS), a maximum suppression of these spontaneous oscillations can be achieved. This analysis suggests that eiDBS in this patient could suppress or amplify spontaneous beta activity in the targeted band (16-22 Hz) with minimum stimulation current as compared with oscillations at other frequency bands.

## Discussion

### Significance and related work

We previously developed the concept of eiDBS using the 1-methyl-4-phenyl-1,2,3,6-tetrahydropyridine (MPTP) nonhuman primate model of PD and showed that amplification or suppression of STN oscillations could be achieved using STN neural responses evoked by stimulation in the GPi[16]. Here, we demonstrate the feasibility of controlling beta band oscillations in the human GPi in real-time by using resonant neural responses evoked by stimulation of the GPi. eiDBS modulates beta oscillations with a stimulation frequency that in average is equal to the mean frequency of the targeted oscillations. Not using high-frequency stimulation is critical to characterizing the role of beta band oscillations in PD, given that high-frequency pulses can improve motor function and also suppress beta band activity. However, it is unclear whether high-frequency stimulation directly reduces beta band oscillations and thereby produces a therapeutic effect, or the reduction in beta band oscillations is unrelated to the therapeutic effect of high-frequency stimulation. Our results inform future studies directed at investigating the causal role of frequency- and location-specific neural activity in the manifestation of specific PD motor and non-motor signs using eiDBS. They are also a step towards developing closed-loop DBS systems that control circuit-wide neurophysiological dynamics associated with brain dysfunction in real-time.

Leveraging the sensing capabilities of directional DBS leads, we showed that the neural sources generating both spontaneous and stimulation-evoked oscillations are likely the same, indicating that 1) eiDBS attains true neural modulation, and 2) observed modulation is not the effect of volume conduction from neural sources located in distinct regions. Furthermore, we showed that patientspecific ER mathematical models combined with LFP recordings can be used to predict the stimulation parameters that maximize the suppression or amplification of spontaneous, frequency-specific neural activity in the human GPi. Therefore, eiDBS can be programmed based on neurophysiological data that is specific to the particular patient.

While we delivered electrical stimulation with precise amplitude and timing (phase), one should note that the rationale behind using phase feedback is different from other approaches intended to induce phase desynchronization across neurons[30,31], modulate depolarization or hyperpolarization of neurons in the proximity of the stimulation site[32], alter short-term plasticity[33], or deliver stimulation based on kinematic variables as tremor[34] or gait[35]. eiDBS continuously overrides the inputs (synaptic-related) of a targeted neuronal population to suppress or amplify spontaneous oscillations via interference created with neural responses evoked by electrical stimulation (low-frequency, synaptic related). The applicability of eiDBS to modulate other brain targets in the human basal ganglia remain to be demonstrated. Nevertheless, our previous pre-clinical studies show that eiDBS delivered in the GPi can be used to modulate frequency-specific neural activity in the STN[16]. Additionally, a previous study with PD patients undergoing DBS surgery indicated that open-loop, isochronal stimulation delivered dorsal to the STN could alter the amplitude of low-frequency oscillations recorded in the STN, whenever consecutive stimulation pulses landed at specific phases of these oscillations[36]. The study presented in [36] suggests that eiDBS could also be employed to modulate STN oscillatory activity in real-time in the human.

### Role of stimulation evoked responses in neural control

Modulation of spontaneous oscillations in the GPi achieved by eiDBS was mediated by neural oscillatory activity evoked by stimulation within the GPi. The pallidal evoked responses resonated at the same frequency (within the beta band) where the spontaneous oscillations resided, suggesting that intrinsic resonant properties of circuits connected to the GPi, thought to play a role in the generation of spontaneous beta band oscillations in PD [37], may also underlie the generation of evoked responses. This resonance in the beta band enabled eiDBS to modulate beta band oscillations with minimal stimulation amplitude as compared with other frequency bands. A reduced stimulation amplitude (and energy) can minimize possible side effects associated with unwanted activation of neuronal pathways and for implantable devices to minimize battery replacements or recharging frequency. While having evoked and spontaneous oscillations with matching frequency can be beneficial, this frequency-matching is not a necessary condition for implementing eiDBS as long as low-frequency, synaptic-related evoked responses are present in the targeted neuronal population.

Although the circuit-wide mechanisms underlying the generation of either spontaneous or stimulation-evoked beta band oscillations in the GPi of PD patients are unknown, these oscillations are likely associated with the activation of multi-synaptic feedback loops in the basal ganglia-thalamocortical network[38–40]. For eiDBS to result in true neural modulation, the ER and spontaneous oscillatory activity need to be generated by the same population of interconnected neurons. The exact location of neural sources within the GPi generating the spontaneous or stimulation-evoked oscillations measured with the directional DBS lead is challenging to estimate using the limited number of LFP channels (i.e., eight) from the lead, restricted spatial coverage of the lead contacts, and uncertain electrical properties of the tissue surrounding the lead contacts. Nevertheless, the tight correlation between the amplitude of spontaneous and stimulation-evoked oscillations across LFP montages created with the directional DBS lead is evidence of these oscillations being generated at the same location and not associated with volume conduction of independent sources at distinct sites. The rationale behind this argument is that two current sources (dipolar or monopolar) located within the same region in a volume conductor generate the same electric potential profile in space (Poisson equation of electrostatics)[28,29].

### Limitations

Our current implementation of eiDBS employs a constant stimulation level to suppress the mean amplitude of the spontaneous oscillations. This approach is suboptimal for suppression of neural activity because it does not account for dynamic changes in the amplitude of spontaneous neural oscillations. Because the stimulation pulse amplitude is set constant, variations in the oscillations amplitude can result in unwanted amplification when the spontaneous oscillations are small, or suboptimal suppression when the spontaneous oscillations are large. Future eiDBS algorithms require instantaneous changes in the stimulation amplitude to precisely suppress spontaneous oscillations in real-time.

Another limitation of the eiDBS algorithm implemented here is that suppression of oscillations in the targeted frequency band resulted in amplification of neural activity in an adjacent band (12-16 Hz). The side-band amplification effect occurs due to phase distortions introduced by the filter at frequencies adjacent to the targeted band, and given that optimal stimulation parameters for one frequency are not optimal for another. This side-band amplification can be a confounding factor when analyzing the effect of suppressing targeted neural activity via eiDBS on brain function. Future eiDBS systems need to track the oscillations’ frequency and reduce filtering-related phase distortions to minimize side-band amplification during the suppression of neural activity.

## Acknowledgments

Research reported in this publication was funded by the Wallin Discovery Fund, the Engdahl Family Foundation, the Kurt B. Seydow Dystonia Foundation, the National Institute of Neurological Disorders and Stroke (P50-NS123109, R01-NS037019), and the University of Minnesota’s MnDRIVE (Minnesota’s Discovery, Research and Innovation Economy) Initiative. We thank Kevin Patino Sosa for building lead adapters and connectors, and Stephanie Alberico and Kevin O’Neill for help during data collection.

## CRediT Authorship Contributions

**David Escobar Sanabria:** Conceptualization, Methodology, Software, Validation, Formal Analysis, Investigation, Resources, Data Curation, Writing - Original Draft, Visualization, Supervision, Project Administration, Funding Acquisition. **Joshua E. Aman:** Resources, Investigation, Data Curation, Writing - Review & Editing, Project Administration. **Valentina Zapata:** Software, Formal Analysis, Data Curation, Visualization. **Luke A. Johnson:** Software, Investigation, Visualization, Writing - Review & Editing. **Hafsa Farooqi:** Software, Data Curation, Visualization. **Jing Wang:** Investigation, Writing - Review & Editing. **Meghan Hill**: Investigation, Writing - Review & Editing. **Remi Patriat:** Data Curation, Writing - Review & Editing. **Kelly Brown:** Project Administration. **Gregory F. Molnar:** Project Administration, Funding Acquisition. **David Darrow:** Investigation, Resources. **Robert McGovern:** Investigation, Resources, Writing - Review & Editing. **Scott E. Cooper:** Resources, Project Administration, Writing - Review & Editing Funding Acquisition. **Noam Harel:** Resources, Funding Acquisition. **Colum D. MacKinnon:** Methodology, Resources, Supervision, Funding Acquisition. **Michael C. Park:** Methodology, Investigation, Resources, Supervision, Funding Acquisition. **Jerrold L. Vitek:** Conceptualization, Resources, Supervision, Writing - Review & Editing, Funding Acquisition, Project administration.

