## Appendix A for "Controlling pallidal oscillations in real-time in Parkinson’s disease using evoked interference deep brain stimulation (eiDBS): proof of concept in the human"

**Evoked response mathematical model.** The following differential equations describe the continuous-time input-output model of the evoked response:

$$\dot{x} = Ax + Bu$$
$$y = Cx,$$

where $x$ is the state vector, $\dot{x}$ is the derivative of $x$ with respect to time, $u$ is the saturated input stimuli (uA), $y$ is the ER output (uV), and $A$, $B$, and $C$ are constant matrices that parameterize the differential equations. These matrices are given as follows.

$$A = \begin{bmatrix} -105.4 & -223.7 & -119.2 & -81.62 & -42.25 \\ 128 & 0 & 0 & 0 & 0 \\ 0 & 128 & 0 & 0 & 0 \\ 0 & 0 & 128 & 0 & 0 \\ 0 & 0 & 0 & 64 & 0 \end{bmatrix}$$

$$B = \begin{bmatrix} 8 \\ 0 \\ 0 \\ 0 \\ 0 \end{bmatrix}$$

$$C = [0 \quad 0 \quad 0 \quad -4.835 \quad 1.013]$$
